# Correlation measures in metagenomic data: the blessing of dimensionality

**DOI:** 10.1101/2024.02.29.582875

**Authors:** Alessandro Fuschi, Alessandra Merlotti, Thi Dong Binh Tran, Hoan Nguyen, George M. Weinstock, Daniel Remondini

**Author notes:** These authors contributed equally to this work (Alessandro Fuschi, Alessandra Merlotti). Deceased (George M. Weinstock).

## Abstract

Microbiome analysis has revolutionized our understanding of various biological processes, spanning human health, epidemiology (including antimicrobial resistance and horizontal gene transfer), as well as environmental and agricultural studies. At the heart of microbiome analysis lies the characterization of microbial communities through the quantification of microbial taxa and their dynamics. In the study of bacterial abundances, it is becoming more relevant to consider their relationship, to embed these data in the framework of network theory, allowing characterization of features like node relevance, pathway and community structure. In this study, we address the primary biases encountered in reconstructing networks through correlation measures, particularly in light of the compositional nature of the data, within-sample diversity, and the presence of a high number of unobserved species. These factors can lead to inaccurate correlation estimates. To tackle these challenges, we employ simulated data to demonstrate how many of these issues can be mitigated by applying typical transformations designed for compositional data. These transformations enable the use of straightforward measures like Pearson’s correlation to correctly identify positive and negative relationships among relative abundances, especially in high-dimensional data, without having any need for further corrections. However, some challenges persist, such as addressing data sparsity, as neglecting this aspect can result in an underestimation of negative correlations.

## Introduction

Techniques based on next-generation sequencing (NGS) can elucidate the complex functioning of natural microbial communities directly in their natural environment. New branches of research have been created such as the study of the human microbiota which showed heterogeneity between different anatomical sites and individual variability [1, 2], or the ability to characterize and monitor the presence of antimicrobial resistance worldwide [3]. Complementing the analyses conducted directly on the abundance of microbiota samples, it can be greatly beneficial to explore a second layer of information represented by the relationships among the observed species. Network theory provides many essential tools to characterize collective properties of the ecology of a natural environment by defining central elements or communities in the system and allowing visualization of these results by exploiting network structural properties [4]. Consequently, the initial step in reconstructing any network involves the identification and quantification of relationships between species, often achieved by assessing correlations or conditional dependencies among each pairwise combination of variables. Independent from the NGS technique used like RNA-seq, 16s or whole genome shotgun, the underlying data are similar, composed of counts of sequencing reads mapped to a large number of references (taxa) and the unifying theoretical framework is their compositional nature [5, 6]. Taxa abundance is determined by the number of read counts, which is affected by sequencing depth and varies from sample to sample.

Typically a sum constraint is imposed over all the samples (1 for probability, 100 for percentage or 10^6^ for part per million) called *L*1 normalization, to remove the effect of sample depth. In this way, data are described as proportions and referred to as compositional data [7, 8]. However, as noted by Pearson at the end of 19th century [9], compositional data can generate spurious correlations between measurements. From a mathematical point of view the data lie on a simplex [8], thus it can be extremely dangerous to use Euclidean metrics for proximity and correlation estimations. These biases on correlation between relative abundances can be significant in some datasets but mild in others [fig:1], and the *diversity* within each sample, called *α*-diversity, (referred to as 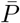, see Materials and Methods) concurs to enforce this bias [10].

**Fig 1.**
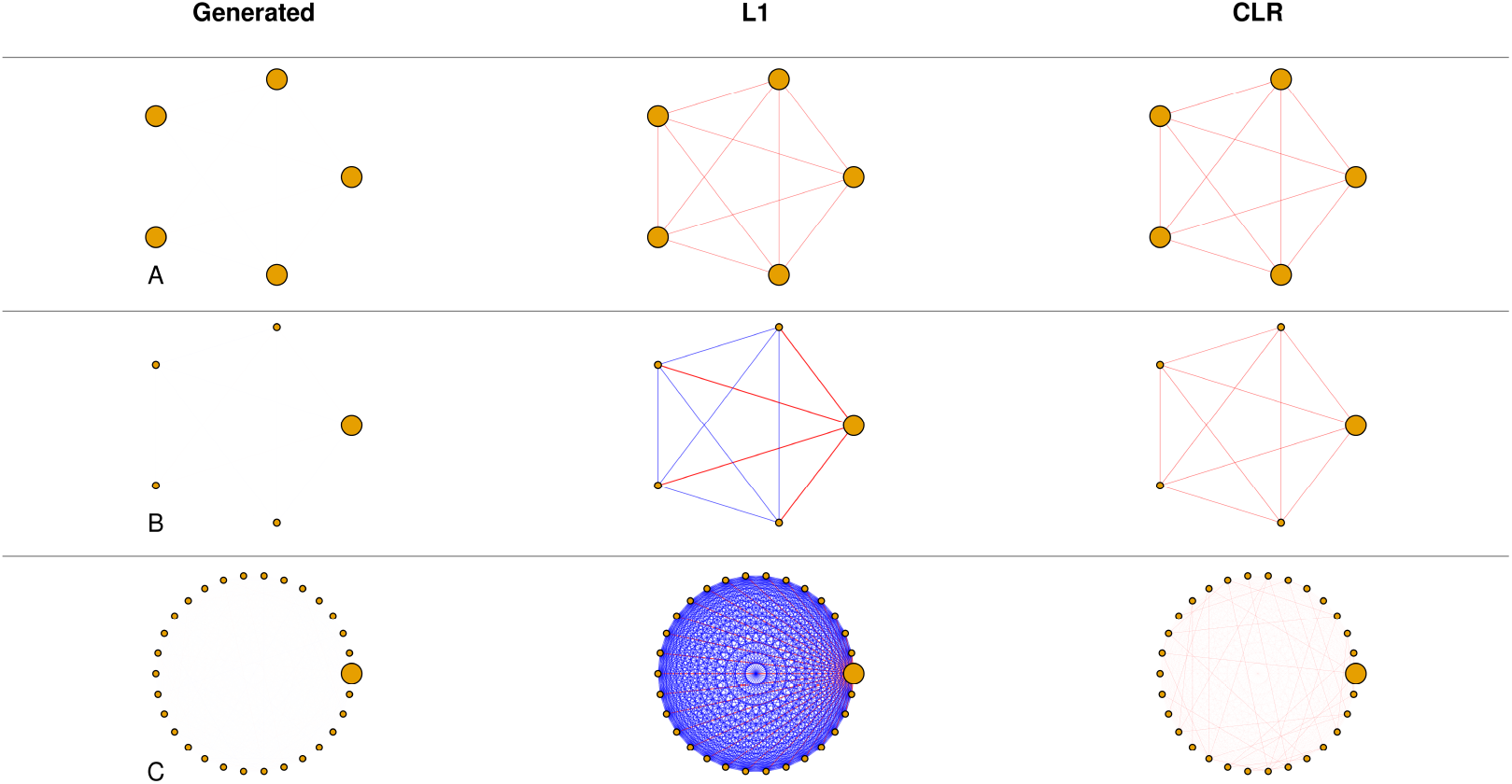
Impact of L1 and CLR Normalizations on Correlation Estimates. Three different cases (A, B, C) are shown, with data generated from uncorrelated multivariate standardized normal distributions sampled 10,000 times, in which data were shifted in order to be positive. Left figures describe the generated data with fixed number of species (dimensionality *D*) and node size proportional to the mean species abundance (*α*-diversity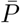); central figures represent Pearson’s correlation as links (red, negative; blue, positive) with width proportional to its value, after L1 data normalization; right figures represent the same situation after CLR data transform. The parameters for the presented cases were: A) *D* = 5 and 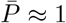, B) *D* = 5 and 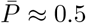, C) *D* = 30 and 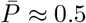. In L1 normalization, biases are strongly associated with dataset diversity and do not decrease with dimensionality, while for CLR normalization these biases decrease with increasing dimensionality and are independent of diversity (see Results Section).

Correlation biases become more pronounced when counts are concentrated in a few taxa. Conversely, when counts are distributed more evenly across samples, these biases tend to decrease. Hence, it is imperative to take into account these compositional effects when reconstructing networks from metagenomic data. Failing to do so may lead to entirely incorrect conclusions [11], endangering the accuracy and reliability of inferred ecological interactions.

To improve correlation estimates on relative abundances, methods such as Sparse Correlations for Compositional data (SparCC) [10], Sparse and Compositionally Robust Inference of Microbial Ecological Networks (SPIEC-EASI) [12], Proportionality for Compositional data (Rho) [13] and many others [14–29] have been developed, almost all making extensive use of the compositional theory introduced by Aitchison [8]. Aitchison provided a family of transformations to handle this type of data, known as log-ratio transformations. The counts of each sample are expressed relative to a reference to enable comparisons, followed by the application of logarithm. One common choice is the centered log-ratio transformation (CLR), where each element is divided by the geometric mean of the sample in a logarithmic scale. This operation is both isomorphic and isometric, preserving distances. However, like L1 normalization, CLR also introduces a sum constraint where the sample sum is fixed to 0. This constraint is equivalent to mapping the counts on a Cartesian hyperplane instead of a simplex, and it also introduces spurious dependencies between variables.

Our work shows that, unlike L1 normalization, the bias introduced by the sum constraint in CLR strongly depends on the dataset dimensionality *D*, or more explicitly it is related to the number of taxa or references [fig:1]. In our study, we not only demonstrate but also quantify these biases, which diminish as the dimensionality increases. In metagenomic contexts, where dimensionality can extend to hundreds or more, the impact of spurious correlations introduced by CLR becomes negligible, making any subsequent step for correlation estimation less critical.

Furthermore, there are additional typical sources of error in the estimation of correlations in metagenomic datasets. Often a large part of taxa in the NGS experiments are under the detection limits of the sequencing techniques, producing very sparse abundance matrices. It’s really common to find datasets where more than 70 − 80% of species are undetected and typically it is assigned the value of 0. The unobserved species are not to be interpreted as the absence of that species but rather as a missing value in which we have no further information. Moreover, non-zero counts exhibit strongly non-normal distributions in non transformed data, with heavy tails that invalidate the assumptions of Pearson’s correlation. The distribution that better describes the real NGS data is still a debated discussion, but in different context the zero-inflated negative binomial distribution (ZINB) is employed [12, 30]. The ZINB distribution can effectively capture the excess of zeros and the dispersion in the data, making it a suitable choice for representing counts in metagenomic datasets, particularly given its discrete nature similar to the counts.

The aim of this manuscript is to explore biases affecting correlation estimates, particularly in the context of compositionality and zero-excess issues commonly encountered in metagenomic datasets. In the absence of a ground truth, we create synthetic datasets across a wide range of conditions, varying dimensionality, diversity, data distribution and sparsity to characterize the biases in correlation estimation. To achieve this, we have developed a model focused on the ‘Normal to Anything’ approach that allows the generation of random variables with arbitrary marginal distributions starting from multivariate normal variables with desired correlation structure.

This work is structured to address three main considerations. The first is the examination of the biases introduced by L1 and CLR transformations in relation to dimensionality and within diversity. This involves a thorough analysis of how these transformations impact data interpretation across various compositional contexts. Importantly, we acknowledge that while CLR is extensively used in metagenomics as a crucial analytical tool, its application is often not accompanied by a deep understanding of its limitations and advantages.

The second consideration corroborates our findings regarding compositional biases arising from L1 and CLR transformations. For this, we compare various recently developed methods on real metagenomic data with the simplest approach of using Pearson correlation on CLR transformed abundances (Pearson+CLR). Our analysis reveals an almost complete overlap in the final results, emphasizing the significance of the CLR transformation.

The third aspect of our research evaluates the role of zero measurements in estimating correlation after minimizing compositional biases through optimal transformation. This involves assessing how zero counts affect the accuracy of correlation measures, thereby providing insights into the appropriate handling of sparse data in metagenomic studies.

## Results

### Compositional biases become negligible with high dimensionality

To comprehend and quantify the compositional biases inherent in Pearson correlation, we conducted a comprehensive comparative analysis. We compared the known correlation structure initially provided as input to the model with the correlation structures obtained after applying L1 and CLR normalizations, while systematically varying the dimensionality *D* and the within dataset diversity 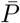 (see Materials and Methods section). In total, we generated 1560 distinct datasets adjusting the dimensionality, ranging from 5 to 200 in steps of 5, and manipulating the within dataset diversity from 0, 025 to 0, 975 in increments of 0, 025, with a tolerance of *±* 0, 005. To isolate the effects of the L1 and CLR transformations, we made deliberate efforts to minimize any known sources of error and chose the simplest experimental conditions to ensure the robustness of our findings. In line with these principles, we consistently conducted the analysis with an uncorrelated covariance structure, and we chose to work with normally distributed variables to avoid potential errors in the Pearson correlations that may result from non-normally distributed data. Furthermore we choose for each experiment *N* = 10000 samples, in order to minimize possible random correlation between variables. Finally, we quantified the biases by calculating the mean absolute error (MAE) on all values of the matrix obtained by subtracting L1- and CLR-normalized correlation matrices, denoted as *R*^*L*1^ and *R*^*CLR*^, to the original correlation matrix R, as follow:

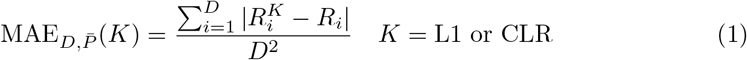

MAE values range within the interval [0, 2], where 0 implies a perfect accordance with the ideal correlation and 2 represents maximum distortion.

The distinct behaviors of the two normalizations are evident, as they introduce different biases on correlation (see Figure 2). Specifically, L1 correlations are primarily influenced by within dataset diversity, with the biases becoming more pronounced as the values within a sample become more heterogeneously distributed. On the other hand, CLR data exhibit biases that are independent of dataset diversity, and these distortions diminish rapidly with increasing dimensionality. Building upon the premise of complete independence of the CLR biases on correlation from dataset diversity, we can estimate this effect by calculating an average over all diversity values 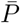. We observe that the error decreases to less than 0.01 for dimensionality values greater than or equal to 100. Thus, we posit that in typical metagenomic scenarios, where the dimensionality often extends into the hundreds, the effects of compositionality are negligible.

**Fig 2.**
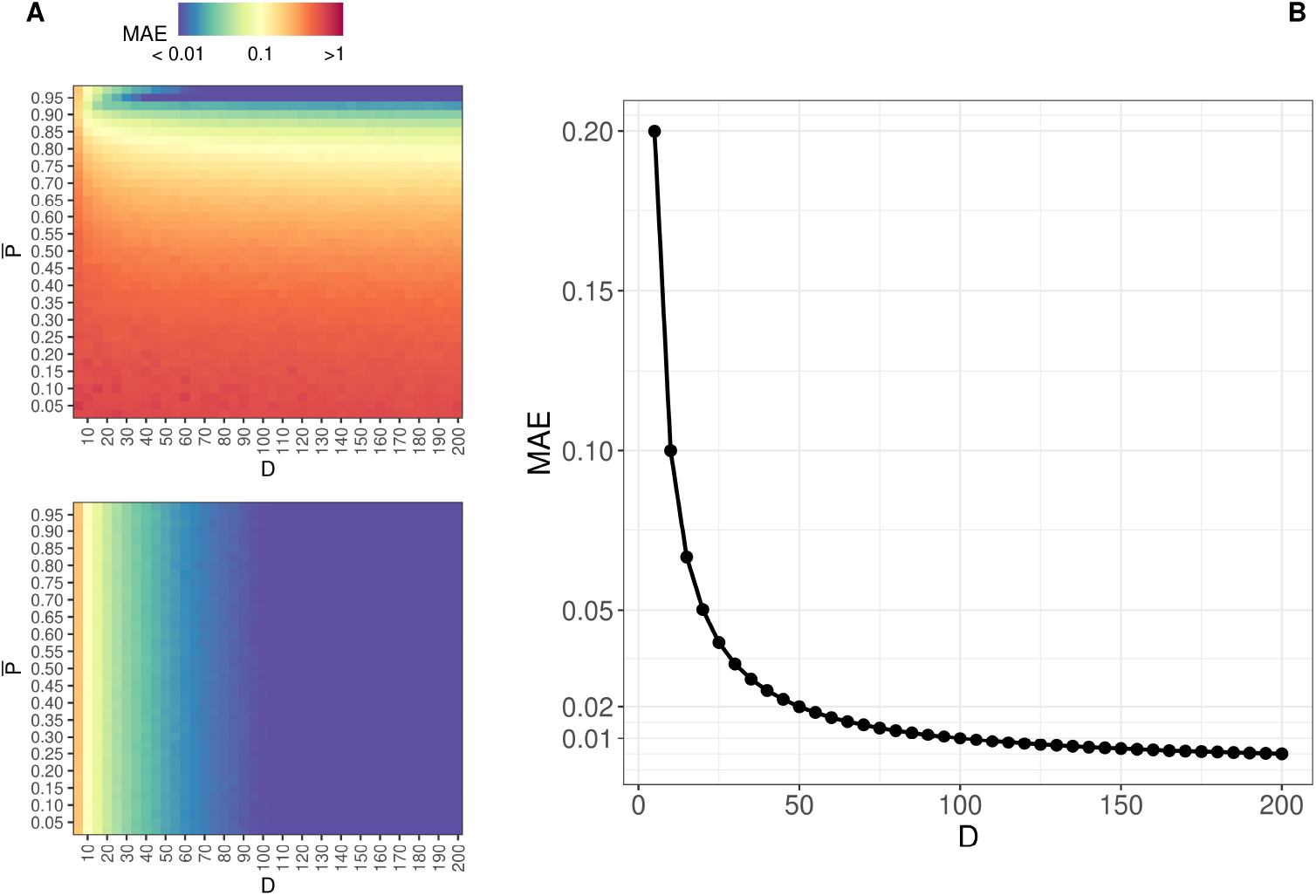
Different behavior of spurious correlations for L1 and CLR transforms : **A)** heatmap of *MAE* as a function of the dataset diversity 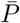and dimensionality *D* for L1 normalization (top) and CLR normalization (bottom) in log_10_ scale. **B)** Scatter plot illustrating the MAE for CLR normalization on correlation as a function of dimensionality.

### Comparison between state of art methods

We opted to compare various computational approaches designed for inferring relationships within compositional datasets, contrasting these with the more straightforward Pearson correlation method applied to CLR-transformed data (Pearson+CLR). This comparison included evaluations of SparCC, the proportionality metric (*ρ*), and the SPIEC-EASI framework utilizing one of its two primary algorithms. All these tools employ similar and comparable concepts, even if developed with different methodologies. For instance, SparCC aims to approximate the Pearson correlation by assuming that the true underlying correlation network is sparse, meaning that highly correlated variables are relatively few compared to the total number. In contrast, *ρ* is based on the similar concept of proportionality as an alternative to traditional correlations, with the goal of mitigating compositional biases.

SPIEC-EASI employs graphical model inference to discern the conditional independence among variables, enhancing its efficacy through iterative evaluations across multiple dataset subsampling. Within SPIEC-EASI exist two inference schemes, we selected the graphical lasso (GLASSO) algorithm for its conceptual alignment with correlation analysis, as it similarly hinges on the covariance structure among the variables. The alternative, the Meinshausen-Bühlmann (MB) algorithm, departs from the correlation-based framework, instead drawing on principles of linear regression for inferring relationships.

All aforementioned methods make extensive use of the compositional theory starting their routine by normalizing data via a log-ratio transformation consistent with Aitchison’s philosophy.

The comparison, conducted on the 51 samples of subject 69-001 in healthy condition from HMP2 (see Materials and Methods) shows an almost complete overlap of the final results, as in Fig. 3. The comparison between the Pearson+CLR with SparCC and Rho is direct, since these three methods produce values between -1 and 1: the scatter plot of the respective correlation values is ≈ 0.99 for both, in very good accordance to a *y* = *x* linear relationship (Fig.3: 1A-1B).

**Fig 3.**
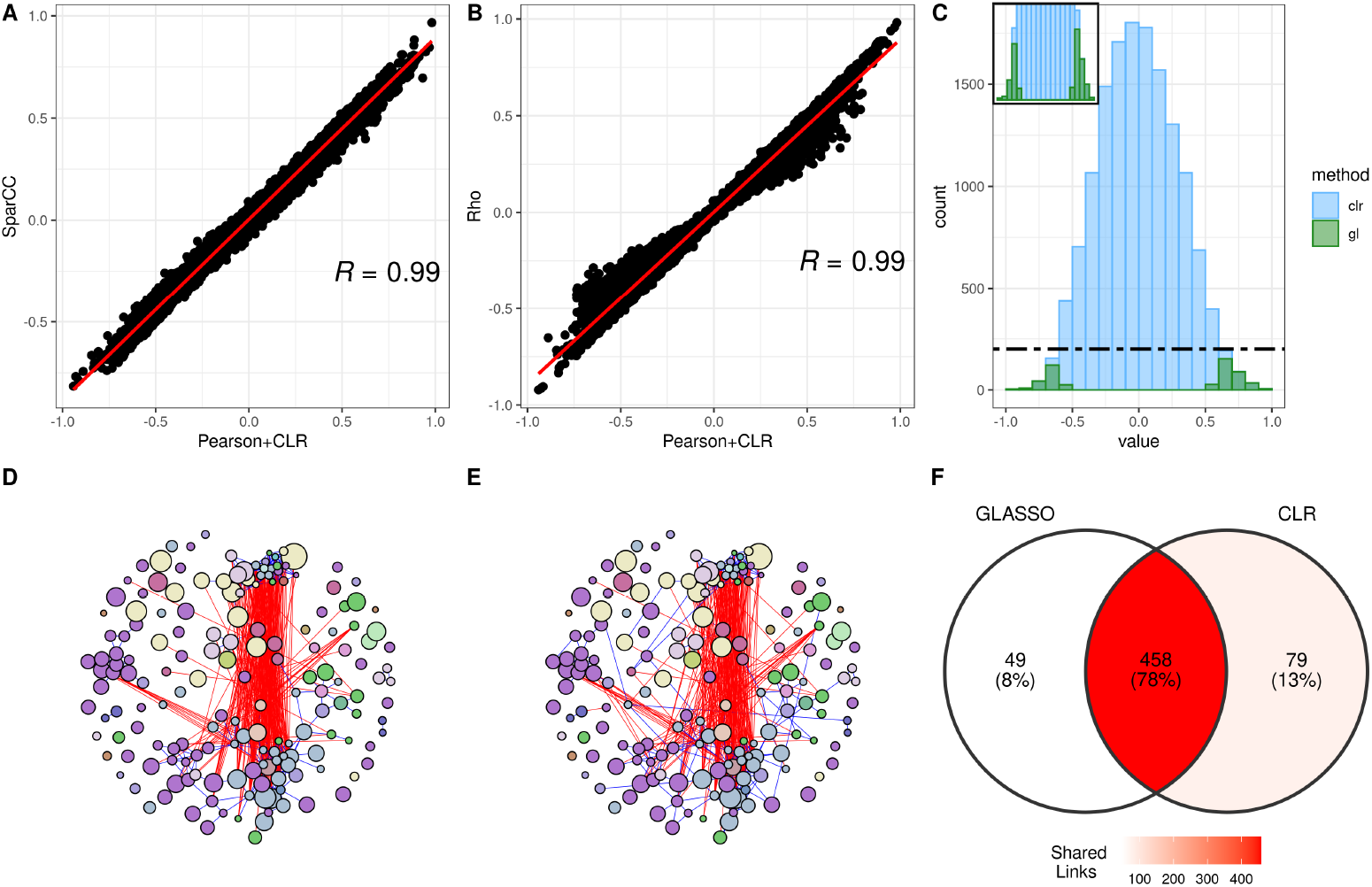
Comparison between state-of-the-art methods and Pearson+CLR on HMP2 data: **A-B)** Scatterplot of the weights associated with pairwise relationships between between the OTUs of Sparcc and *ρ* compared to Pearson+CLR. **C)** Significant links obtained through the SPIEC-GLASSO mapped to the Pearson+CLR correlation histogram. **D)** Reconstructed network from Pearson+CLR method with Bonferroni-corrected p-value threshold set at 0.05; **E)** Reconstructed network using the SPIEC-EASI GLASSO method with a stability threshold set to 0.05; we used the same vertices layout as for Pearson+CLR network. **F)** Venn diagram of the shared links between SPIEC-EASI GLASSO and Pearson+CLR networks.

Since SPIEC-EASI produces a binary output in terms of conditional independence between each pair of variables, we consider the histogram of Pearson+CLR values, and overlap bins corresponding to couples of variables significantly associated through SPIEC-EASI by imposing a threshold on overall stability equal to 5%. Most of the significant links for SPIEC-EASI are associated to high absolute values of Pearson+CLR (Fig.3: 1C). This analysis shows that significant SPIEC-EASI associations predominantly correspond to high absolute Pearson+CLR values.

Further comparison were performed between the networks inferred by the SPIEC-EASI GLASSO and Pearson+CLR through thresholding, considering as links the correlations with *p <* 0.05 after Bonferroni correction for multiple testing. Approximately 78% of the edges were common between the inferred networks (Fig 3: 2C), and a visual inspection of the network representations indicates that their collective properties are nearly identical (Fig 3:2A-2B).

In practical applications, despite the heterogeneity of their underlying methodologies, the considered methods converge towards equivalent outcomes. This observation underscores the central role of the Centered Log-Ratio in all considered algorithms, that is sufficient to minimize spurious correlations within high-dimensional contexts.

Particularly in metagenomic studies, where the dimensionality often extends into the hundreds, the necessity for additional corrective measures appears redundant.

### Data sparsity remains a limitation

In this section we focus on the error on estimating correlation as a function of the ratio of zero values in the samples, similar to real-world scenarios. To achieve this, we have implemented a zero-inflated negative binomial distribution as the target distribution within our modeling framework based on NorTA approach (see Materials and Methods section). This distribution was selected to accurately capture the frequent occurrence of zero counts and the asymmetrical distributions seen in real data.

In the preceding section, we discussed measures taken to minimize the impact of spurious correlations introduced by the CLR transformation. To achieve this, we standardized the dimensionality (*D*) of all generated datasets to 200, a choice informed by its effectiveness in ensuring that correlation errors remain consistently below the threshold of 0.01. Even in this analysis we fixed the number of observations (*N*) to 10^4^ to reduce errors within the estimated correlation matrix. Furthermore, we only took the CLR into consideration for the analysis given that the L1 in real situations, with more heterogeneously distributed data, is impractical as seen in the previous section.

We generate data that closely resemble real-world observations deriving the parameters *munb, size* and *ϕ* of the zero-inflated negative binomial distribution from the actual distributions of the OTUs of subject 69-001 in the HMP2 dataset, using the *fitdist* function from the R package *SpiecEasi* [31]. Each taxon was then generated using random parameters falling within the range of the first and ninth deciles of the previously fitted ZINB parameters, distributed according to their empirical distribution using the *quantile* function of base R.

To quantify the error, we consider the absolute difference between the initial data correlation matrix *R* and the correlation on the same data transformed through NorTA approach and CLR, *R*_*CLR*_ with nonzero correlation only between two taxa labeled *I* and *J*. We build the correlation matrix specifically by varying only the value between I and J, labelled as *r*, from − 0.9 to 0.9 in steps of 0.05, leaving all the others 198 taxa uncorrelated. In practice, all the other taxa other than I,J only contribute to reduce the biases introduced by the CLR transformation. Moreover, we varied the ratio of zero counts (*ϕ*_*I*_ and *ϕ*_*J*_) of their respective marginal distributions from 0 to 0.95 in increments of 0.025. This process enables us to track the correlation error between taxa *I* and *J* across different levels of sparsity and correlation (err_*ϕ*,*r*_).

This process was repeated 100 times for every combination of *ϕ* and *r*, and MAE was calculated as follows (see Fig. 4.A):

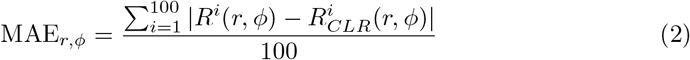

**Fig 4.**
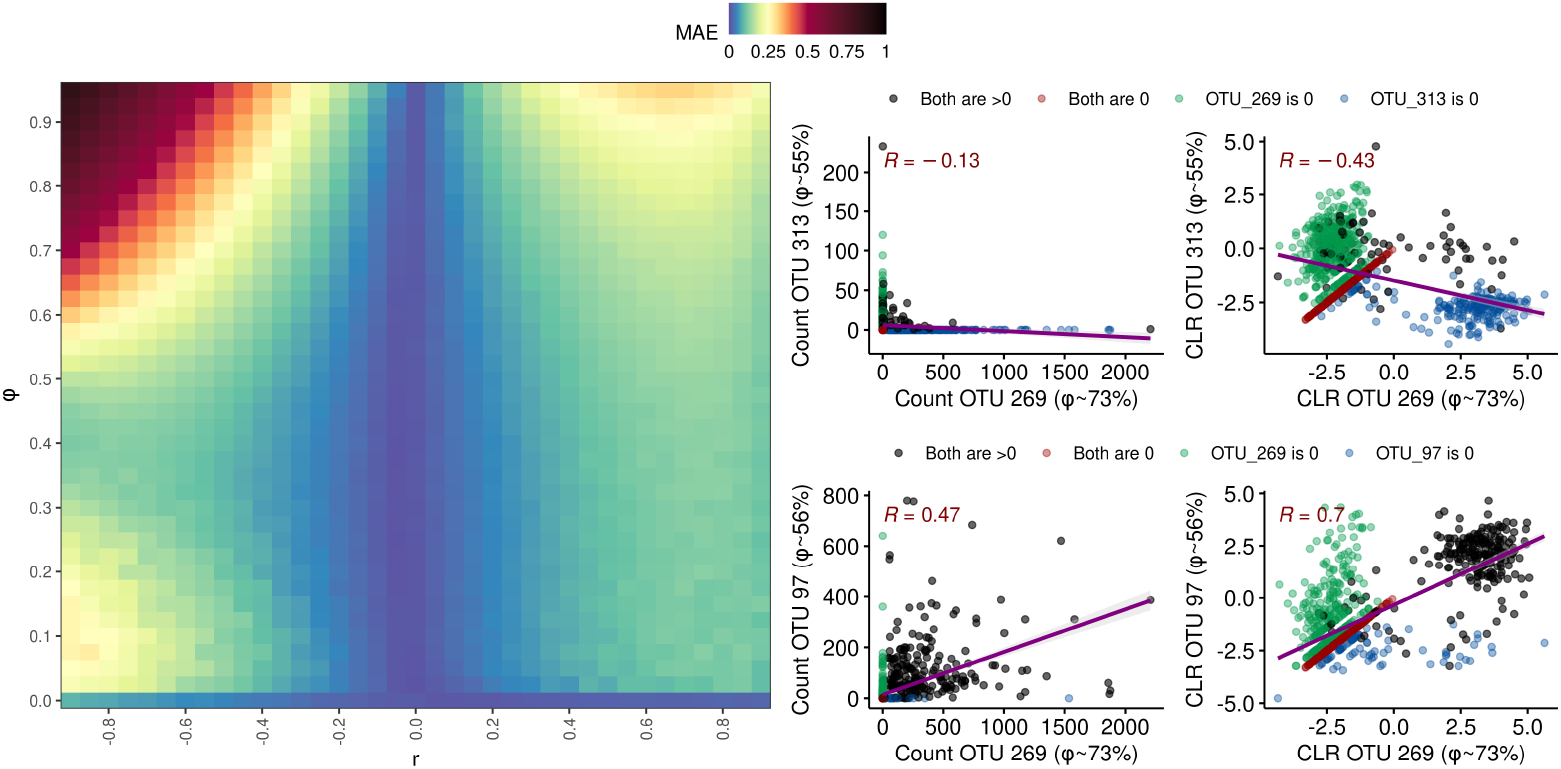
Impact of Sparsity on Correlation Coefficients in CLR-Transformed Data: **A)** heatmap depicting MAE of correlation coefficients across different values of sparsity Φ and correlation values *r*. **B)** effect of CLR transform on negative (top subplots) and positive (bottom subplots) correlation between selected pair of OTUs from HMP2 dataset, with a significant presence of zero counts, before (left subplots) and after (right subplots) CLR transform.

An important aspect to emphasize in our methodology is the deliberate decision to randomly generate parameters for each ZINB distribution. This approach was intended to observe the correlation phenomenon in a manner that is as independent as possible from any specific data distribution, ensuring that our findings are not biased by particular distributional characteristics of the data. The pseudo-code below summarizes our methodology:

~~~
// Fit ZINB Model parameters using OTUs from HMP2
params_ZINB_HMP2=fitZINBParameters(OTUs_HMP2);
// Perform 100 iterations of simulation
for (iteration in 1:100) {
    // generate ZINB random parameters using the ecdf of the real
    // distributions of the ZINB parameters
    random_params_ZINB=randomZINBParameters(params_ZINB_HMP2);
    // Generate synthetic dataset with D=200
    syntheticData = generateSyntheticDataset(D=200, par=random_params_ZINB
   // Loop over varying levels of sparsity (phi) and correlation (r)
   for (phi in seq(0, 0.95, by=0.025)) {
       for (r in seq(-0.9, 0.9, by=0.05)) {
           // Modify variables I and J in the dataset
           modifyVariables(syntheticData, I, J, phi, r);
          // Record error for current sparsity and correlation
          err_phi_r = recordError(syntheticData, I, J);
       }
   }
}
// Calculate the Mean Absolute Error (MAE) for each phi and r
// over the 100 iterations
calculateMAE(err_phi_r);
~~~

MAE significantly differs between positive and negative correlations, as clearly illustrated in (Fig. 4-A). While the error generally grows with an increasing number of zeros, this effect is particularly marked for taxa with negative correlations, as observed in the upper left section of the figure. Additionally, it is noteworthy that when variables are uncorrelated, the presence of zeros does not significantly impact the results.

An important aspect in the application of the CLR transformation is the number of zero counts, that requires the introduction of pseudo-counts to avoid logarithm divergence. This is illustrated in Fig. 4-B, using data from the HMP2 dataset, where we consider two pairs of OTUs with a high percentage of zeros and opposite sign of the correlation values. When examining negatively correlated variables in metagenomic studies, most of the nonzero values of one variable are matched with the pseudo-counts of the other. Such a pattern leads to a flattening on the *x, y* axes of the two OTU scatterplot, producing a hyperbolic-like pattern (Fig. 4-B top left) that tends to underestimate the value of negative correlation. We show that CLR significantly increases the negative correlation value mitigating this phenomenon, also in case of positively correlated OTUs (Fig. 4.B bottom).

## Discussion

The network analysis framework is a robust tool for enhancing our comprehension of metagenomic studies, enabling us to unravel the intricate dynamics of microbial ecosystems. Although network reconstruction from second-order statistics such as correlation offers a straightforward methodology, the compositional nature of metagenomic data presents unique analytical challenges that require specialized techniques. Our study conducts a detailed investigation into the potential biases that affect the accuracy of correlation measures, considering factors such as dimensionality, diversity, and sparsity of datasets,characteristics commonly associated with metagenomics data of any type.

Our analysis is focused on the effect of the Centered Log-Ratio (CLR) transformation when applied to compositional data. We discovered that the spurious correlations introduced by the CLR transformation decrease as a function of sample dimensionality. This contrasts with the L1 transformation, where spurious correlations are mainly influenced by the within-diversity of the dataset and do not decrease with sample dimensionality. Given the high dimensionality that characterizes metagenomic datasets— in the order of hundreds or more OTUs or taxa—the spurious correlations associated with CLR become thus negligible. The CLR transformation is also adequate to rectify the effect of diversity for sufficiently high-dimensionality data (in the order of hundreds) without additional adjustments, at difference with L1 transform for which high diversity remains an issue. We underscore the pivotal importance of the CLR transformation as a foundational step for metagenomic studies, streamlining the processing steps while ensuring data integrity.

To validate the role of the CLR transformation in compositional data analysis, we conducted a comparative study using various state-of-the-art algorithms specifically designed to estimate associations in metagenomic datasets. Our findings indicate a striking convergence of SparCC, *ρ*, and SPIEC-EASI GLASSO methods for correlation estimation towards Pearson’s correlation on CLR-transformed data. This convergence suggests that the log-ratio transformation is the critical normalizing step across all methods, effectively neutralizing the compositional bias inherent to the data.

However, we must also acknowledge the substantial impact of dataset sparsity on correlation measures: the large number of zero counts associated with low-abundance taxa can significantly distort correlations, more severely affecting negative correlations. While CLR mitigates this distortions, the proportion of zero counts is the crucial parameter: the larger the zero count ratio, the larger the distortion. It is thus impossible to entirely eliminate the bias introduced by zero counts, unless eliminating any information about very rare species. A compromise must thus be found between minimizing correlation distortions and retaining low-abundance species in the analysis. This trade-off is fundamental for ensuring the accuracy and comprehensiveness of metagenomic data interpretation as a function of the study design.

## Materials and Methods

### Within Dataset Diversity 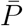

The within diversity of a dataset 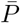 is defined as the mean value over all the samples of the Pielou index [32], which is the Shannon entropy normalized to 1 with respect to the dimension. Given a dataset **X** ∈ *𝒩*^*N*,*D*^ composed of *N* distinct samples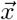 of dimension *D*:

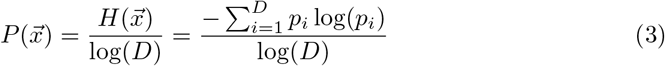

with 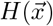 the Shannon entropy, *p*_*i*_ corresponding to the i-th taxa relative abundance in the sample. Finally, the diversity of a dataset 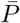 is calculated as:

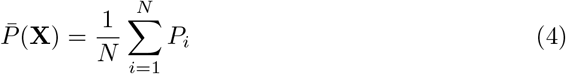

### Generation of Gaussian Data for Characterization of L1 and CLR Correlation Biases

Our examination of the biases introduced by L1 and CLR transformations began with the creation of synthetic datasets modeled on Gaussian distributions. This methodology was specifically crafted to underscore the compositional biases inherent in metagenomic datasets, with a concentrated focus on dimensionality (*D*) and within-sample diversity 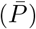—elements that are fundamentally tied to the compositional nature of the data. Our objective was to isolate and examine biases arising specifically from these compositional attributes, recognizing their direct impact on correlation analysis. While we acknowledge that sparsity and non-Gaussian distribution patterns also affect correlation metrics, these elements are secondary in the context of compositional data analysis. They were thus delineated outside of this study’s primary scope and are addressed in a subsequent section.

Utilizing the *mvtnorm* R package [33], we constructed a matrix of variables following a multivariate Gaussian distribution. In this matrix, the dimension *D* corresponds to the variables (or taxa), and *N* signifies the number of observations or samples, all governed by a predefined correlation matrix. To enable the calculation of the Pielou index without modifying the correlation structure, all generated values were shifted to be positive.

To tune the within dataset diversity of the generated Gaussian data, a simply but functional strategy was employed: applying a multiplicative factor to one selected variable from the Gaussian-generated dataset. This deliberate manipulation skewed the distribution towards this variable, thus altering the dataset’s diversity 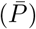 without distorting the established correlation structure.

Following this adjustment for within dataset diversity, both L1 and CLR transformations were applied to the synthetic datasets. We then extracted the correlation matrices from these transformed datasets to analyze the biases each normalization method introduced.

### Realistic Synthetic Data Generation for Sparsity Biases Characterization on Correlation Measurement

To generate realistic artificial data with specified characteristics such as dimensionality (*D*), correlation structure (*R*), and sparsity (Φ), we have extensively used the ‘Normal to Anything’ (NorTA) paradigm. This framework is capable of producing an arbitrary multivariate distribution that conforms to a pre-established correlation structure *R*, drawing upon the principles of copula functions theory [34]. Essentially, the NorTA method allows for the transformation of normally distributed data into any desired distribution while preserving the original correlation structure. The core principle of the NorTA approach involves two main steps: Firstly, generating a multivariate normal dataset with the desired correlation structure, and secondly, transforming this dataset to have the targeted distribution while maintaining the predetermined correlations. The transformation is mathematically represented as follows:

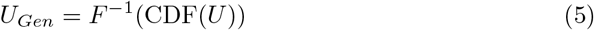

In this equation, *U* represents the multivariate normal data, CDF is the cumulative distribution function of the normal distribution, *F*^−1^ is the inverse CDF (quantile function) of the target distribution, and *U*_*Gen*_ is the transformed data with the desired distribution and correlation structure.

We have already defined key parameters of the generated dataset, indeed the dimensionality (*D*) and the correlation structure (*R*) are trivially integrated within the NorTA framework. However, the delineation of dataset sparsity (Φ) is a less obvious aspect, it is determined by the selection of the marginal distribution *ρ*. To introduce sparsity we have to appeal to the zero-inflated or the hurdle versions of conventional distributions. These modified distributions include an additional parameter, commonly denoted as *ϕ*, which regulates the proportion of zero-valued data. Thus, the level of sparsity within the final dataset Φ depends from the *ϕ*_*i*_ parameters designated for each marginal distribution.

Finally, we perform L1 and CLR transformations on the tuned dataset *U*_*Gen*_, yielding *U*_*L*1_ and *U*_*CLR*_, respectively, each with their corresponding correlation matrices *R*_*L*1_ and *R*_*CLR*_. The central goal of our model is to assess how these transformations impact the correlation matrices in comparison to the original matrix *R*, and not respect the empirical matrix from *U*_*Gen*_. Specifically, we aim also to evaluate the CLR transformation’s efficacy in addressing the skewness and normalizing data with heavy-tailed distributions through logarithmic scaling.

Since the CLR transformation is not defined for zero values, we replaced them with a value corresponding to the 65% of the sample detection limit, in order to minimize the distortion in the covariance structure, as in [35, 36].

### HMP2 16S Human Gut Data

We utilized the Human Microbiome Project’s second iteration (HMP2) dataset, which encompasses operational taxonomic unit (OTU) counts and taxonomic classifications from a longitudinal study on the microbiomes of healthy and prediabetic individuals over a period of up to four years [40]. The complete dataset includes 1122 samples encompassing 1953 OTUs derived from 96 subjects. Each sample is accompanied by metadata indicating the health status of the corresponding subject. To enhance the homogeneity of the dataset for our analysis, we narrowed the focus to a single subject coded as 69-001, who is classified as healthy and has contributed 51 samples. To refine the dataset further, we applied a filtering process based on OTU prevalence and median values of the abundances. Specifically, we retained OTUs with non-zero values in *>* 33% of the samples and a median value of non-zero counts ≥ 5. This stringent selection criterion was designed to eliminate the rarest OTUs and focus on those with a consistent presence across the samples, thereby facilitating a more robust subsequent analysis.

## Data and Code Availability

For free access to all the code and data utilized, please visit the following URL: https://github.com/Fuschi/Correlation-Biases-on-Metagenomics-Data - GitHub Repository. This repository contains comprehensive resources for replicating the analyses based on R base [37], *VGAM* [38], *mvtnorm* [33], and *igraph* [39].

## Acknowledgments

D. R. and A. F. acknowledge EU H2020 “VEO - Versatile Emerging infectious disease Observatory” Project n. 874735 and EU H2020 ERA-HDHL “SYSTEMIC - An integrated approach to the challenge of sustainable food systems” n. 696295.

The authors would like to thank G.W. for inspiring this work through fruitful discussions and joint work. His loss is a big miss for all of us.

